# Competition for resources can reshape the evolutionary properties of spatial structure

**DOI:** 10.1101/2024.04.13.589370

**Authors:** Anush Devadhasan, Oren Kolodny, Oana Carja

**Affiliations:** Computational Biology Department, School of Computer Science, Carnegie Mellon University, Pittsburgh, PA, USA; Department of Ecology, Evolution, and Behavior, E. Silberman Institute of Life Sciences, The Hebrew University of Jerusalem

**Keywords:** population structure, evolutionary graph theory, eco-evolutionary dynamics, specialist populations, generalist populations, probabilities of fixation, time to fixation, graph-structured populations

## Abstract

Many evolving ecosystems have spatial structures that can be conceptualized as networks, with nodes representing individuals or homogeneous subpopulations and links the patterns of interaction and replacement between them. Prior models of evolution on networks do not take ecological niche differences and eco-evolutionary interplay into account. Here, we combine a resource competition model with evolutionary graph theory to study how heterogeneous topological structure shapes evolutionary dynamics under global frequency-dependent ecological interactions. We find that the addition of ecological competition for resources can produce a reversal of roles between amplifier and suppressor networks for deleterious mutants entering the population. Moreover, we show that this effect is a non-linear function of ecological niche overlap and discuss intuition for the observed dynamics using simulations and analytical approximations.

## Introduction

Coexistence is a hallmark of many natural systems and the maintenance of biodiversity is a central theme in ecology and conservation biology (Wu et al., 2021; Patel et al., 2014). The mechanisms of coexistence can often be traced to differences in resource utilization that circumvent competitive exclusion and allow longterm stable maintenance of two or more ecotypes in the population (Chesson, 2000b; Barabás et al., 2018; Chesson, 2000a; Chase and Leibold, 2009; MacArthur, 1970). Until recently, most models assumed the time scales of ecology and evolution to be non-overlapping, and rarely considered the role of constitutively beneficial fitness mutations and stochastic evolutionary dynamics in perturbing these ecological equilibria (Sigmund and Holt, 2021; Vallespir Lowery and Ursell, 2019). However, it has become increasingly clear that rapid adaptation can occur on the timescale of ecological niche diversification, for example through the stochastic accumulation of adaptive mutations within ecotype clades, resulting in fixation of particular lineages and the eventual loss of coexistence (Herron and Doebeli, 2013; Traverse et al., 2013; Good et al., 2017, 2018).

While it is increasingly theoretically and empirically recognized that ecological interactions can shape evolutionary dynamics, studies usually assume well-mixed populations and no spatial heterogeneity (Good et al., 2018). However, organisms live in spatial communities (Torres-Pulliza et al., 2020) and empirical work is reinforcing the importance of heterogeneous networks of interaction and replacement in shaping the genetic makeup of eco-evolutionary systems. Spatial and temporal gradients have been shown to interact to shape the dynamics of plant–pollinator networks (Marcacci et al., 2023) and the complex branching topology of riverine ecosystems has been shown to be critical for biodiversity maintenance (Terui et al., 2021). Wastewater networks have been shown to harbor complex spatial ecosystems of critical importance for human health due to their potential to harbor antibiotic resistance gene-carrying bacteria (Medina et al., 2020). Microbial spread and colonization of the nasal turbinates has been shown to be patchy with single bacterial cells seeding the surface of epithelia at seemingly random locations (Swidsinski et al., 2007) and we are just beginning to understand how surface topography, i.e. the network structure of contact between different locations, determines the importance of spatial pattern and scale in shaping evolutionary dynamics in commensal microbial communities within our bodies (Proctor et al., 2020; Proctor and Relman, 2017).

Prior models that investigate the role of spatial structure in shaping eco-evolutionary dynamics have used a meta-community framework to understand how habitat fragmentation and destruction reshape eco-evolutionary feedback loops (Mullon and Lehmann, 2018; Legrand et al., 2017). In parallel, using the mathematical proxy of a network for representing complex structures of interaction and replacement, evolutionary graph theory (EGT) has been developed as a modeling framework to study evolution on network-based populations (Lieberman et al., 2005).

Initial results have highlighted that tuning spatial heterogeneity beyond what is possible with regular deme and island-based models can significantly extend the range of possible evolutionary outcomes for a population (Lieberman et al., 2005; Ohtsuki et al., 2006; Allen et al., 2017; Kuo et al., 2021; Kuo and Carja, 2024). The evolutionary properties of a graph can be understood and predicted by how they change the rate of evolution of a new mutant appearing on the network, compared to a well-mixed population. These properties can be summarized by two parameters, the amplification and acceleration factors, which characterize the change in the probability and time to fixation of the new mutant, compared to those in a well-mixed or regular lattice-based population (Kuo et al., 2021; Kuo and Carja, 2024). For a new mutant with a fixed selection coefficient *s* entering the population, evolutionary graph theory classifies population topologies as one of two types: amplifiers or suppressors of selection. An amplifier of selection boosts the selective benefit of new beneficial mutants, compared to a well-mixed population (or equivalently, the complete graph), and lowers it if the mutant is deleterious, with the switch occurring precisely at neutrality, *Ns* = 0. The opposite is true for suppressor topologies (Lieberman et al., 2005).

Here we discuss how this simple, but powerful classification breaks down in significant and unintuitive ways when we introduce global frequency-dependent ecological interactions. While the framework of graph-structured populations has been previously used in ecological models to model spatially heterogeneous resource distributions beyond simple structures of dispersal, prior models do not explicitly incorporate frequency-dependent ecological diversification pressures (Kaveh et al., 2020; Nemati et al., 2023). In this paper we develop and investigate a model that combines the EGT framework with a resource competition model and study how the existence of ecological niche differences reshapes probabilities of fixation of new mutants in network-structured populations. In the limiting case of complete niche overlap, our model recovers the constant directional selection case. With decreasing niche overlap however, we find that the interplay between frequency-dependent ecological selection and frequency-independent directional selection makes spatial structures that are amplifiers of selection increase probabilities of fixation for weakly deleterious mutants, in contrast to previous results in EGT. A similar reversal of roles effect is observed for suppressors of selection.

Our results demonstrate that the complex topology of many real world evolving ecosystems, ranging from complex wastewater networks that harbor antibiotic-resistant microbes to riverine ecosystems with intricate branching structures, in interplay with the structure of ecological interaction, can critically reshape their genetic makeup. Our work develops an initial framework to understand the effect of spatial structure on eco-evolutionary dynamics as well as inform on how to spatially engineer these types of ecosystems to select for evolutionary outcomes of interest.

## Model

To study the role of heterogenous population structure in shaping eco-evolutionary dynamics, we use a resource-competition model in which individuals with different metabolic strategies compete for an assortment of externally supplied resources. This resource-based model has the benefit of capturing the key evolutionary and ecological features observed in natural and experimental populations (Herron and Doebeli, 2013; Lang et al., 2009), while remaining analytically tractable across a large range of parameters (Posfai et al., 2017). We consider the simplest nontrivial scenario of two ecotypes evolving in an environment with just two resources. This two-resource case is sufficient to explore the key qualitative differences introduced by ecological interactions, while allowing for analytical tractability. We assume a population of fixed size *N* and analyze the probability of fixation of a single mutant ecotype (mut) invading an initially homogenous wild-type (wt) population.

To represent heterogeneous population spatial structure, we use unweighted and undirected graphs, where each node in the graph represents one individual in the population and edges are proxies for the local pattern of replacement and substitution. To model population dynamics we use a stochastic discrete-time Moran model with Birth-death (Bd) updating. The population is initialized with exactly one mutant, placed on a random node of the population network. At each discrete time step, one individual is sampled for reproduction proportional to fitness. The offspring then replaces a randomly selected neighbor of the parent node. This is the essential difference that allows us to study the role of local population structure, in comparison to a well-mixed population: in a well-mixed population, a node from the entire network would be randomly selected for death.

While the population is assumed to have a network-based structure of reproduction and replacement, the resource availability is spatially uniform: each individual in the population has equal access to the resources, driving the global frequency-dependent dynamics on the whole network. Therefore, our model applies to biological examples where resources can be assumed well-mixed, i.e. bacteria residing in wastewater network biofilms that rely on resources flowing through wastewater, plant communities that rely on resources flowing through rivers, plants relying on pollinators that diffuse freely through the air. We also present a relaxation of the assumption of spatially homogeneous resources and show that our results hold in **Supplementary Figure S5**.

The two resources are externally supplied at a constant rate *S*. Each ecotype *σ* (mut or wt) is defined by its resource uptake metabolic strategy, the proportion of its metabolic resources allocated to the consumption of each of the two nutrients, *α*_*σ*_ = (*α*_*σ*1_, *α*_*σ*2_). The rate of uptake of each nutrient is linear in its concentration and the nutrient concentration dynamics of resource *i, c*_*i*_, follows

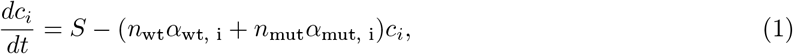

where *n*_*σ*_ is the number of individuals of ecotype *σ* in the population at time *t*. We assume a negligible degradation rate.

Finally, to incorporate evolutionary dynamics, we introduce a pure fitness difference parameter, orthogonal to the ecological dimension of resource utilization, that can capture either a potential change in the metabolic efficiency of the new mutant lineage or reflect phenotypic changes not directly involved in niche diversification. Without loss of generality, we assume a wild-type pure fitness *s*_*wt*_ = 1 and *s*_*mut*_ = (1 + *s*) for the mutant.

To focus the analysis on how a switch in the preferred resource to one that is under-utilized by the wild-type population affects the evolution of the mutant ecotype, we assume *α*_wt_ = (1−*α, α*) and *α*_mut_ = (*α*, 1−*α*). In other words, the mutant ecotype *mut* switches the preferred resource of the individual, while leaving the relative uptake rates of the preferred and secondary resources intact (but see **Supplementary Figure S4** for a relaxation of the assumption of symmetric uptake rates). In this symmetric model, both species have a fixed metabolic budget, i.e. *α*_*σ*,1_ + *α*_*σ*,2_ = 1. This ensures that the pure fitness difference between wild-type and mutant is driven only by differences in metabolic efficiency.

This also allows us to use the parameter *α* as sole proxy of the strength of ecological selection arising from niche overlap. We will therefore refer to this component of fitness that relates to niche partitioning as the “ecological” component of fitness. Parameter *α* ranges from a minimum value of 0.5, which corresponds to no ecological selection dynamics (’generalist’ phenotypes), to a maximum value of 1, corresponding to a regime of completely orthogonal niches, where each ecotype consumes only one of the two resources (’specialist’ ecotypes). Taking into account the two components of fitness, the overall fitness of ecotype *σ* is given by

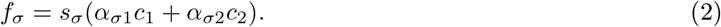

Ecological selection and niche differences between the two ecotypes introduce negative frequency dependence that pushes the lineages towards an interior equilibrium of coexistence, which depends on the pure fitness difference *s*. This equilibrium is unstable however and, due to genetic drift, the only stable equilibria are the two absorption points, fixation or loss of the mutant allele.

We develop analytic approaches and use Monte-Carlo simulations with at least 10^6^ runs to estimate the mutant’s fixation probability and conditional fixation time. We systematically explore the space of possible network topologies by generating structures from a variety of network families, including random spatial geometric graphs, preferential attachment, bipartite and small-world graphs (see **Materials and Methods** for a description of the network families used in this study). To further analyze the space of topologies within graph families, we utilize the concept of the dk-distribution (Mahadevan et al., 2006) and systematically tune different network properties, independently of one another, across levels of organization. Specifically, we use simulated annealing with edge-swap operations to sample from the full range of a network’s dk-distribution, while keeping all lower levels constant (Kuo et al., 2021).

## Results

Previous work in evolutionary graph theory has shown that, for single, pure fitness mutations, the evolutionary role of complex spatial structure can be quantified by analyzing two essential network properties: the network amplification factor, which shapes probabilities of fixation compared to well-mixed populations (Kuo et al., 2021) and the network acceleration factor, which shapes the time to fixation for new mutations in the population (Kuo and Carja, 2024). In particular, the network amplification factor, *a*_*Bd*_, quantifies how to rescale the selection coefficient *s* for the well-mixed model to obtain the same probability of fixation as an allele with selection coefficient *s* on a network population. In other words, a mutation with a selective benefit *s* in a graph population, would have a probability of fixation corresponding to an equivalent selection coefficient *a*_*Bd*_*s* in a well-mixed population. If *a*_*Bd*_ *>* 1, the graph amplifies selection, if *a*_*Bd*_ *<* 1 the graph suppresses selection and if *a*_*Bd*_ = 1 the graph does not change probabilities of fixation compared to the well-mixed. Example dynamics for an amplifier under a model of no ecological selection is presented in **Figure 1A** (black lines).

**Figure 1.**
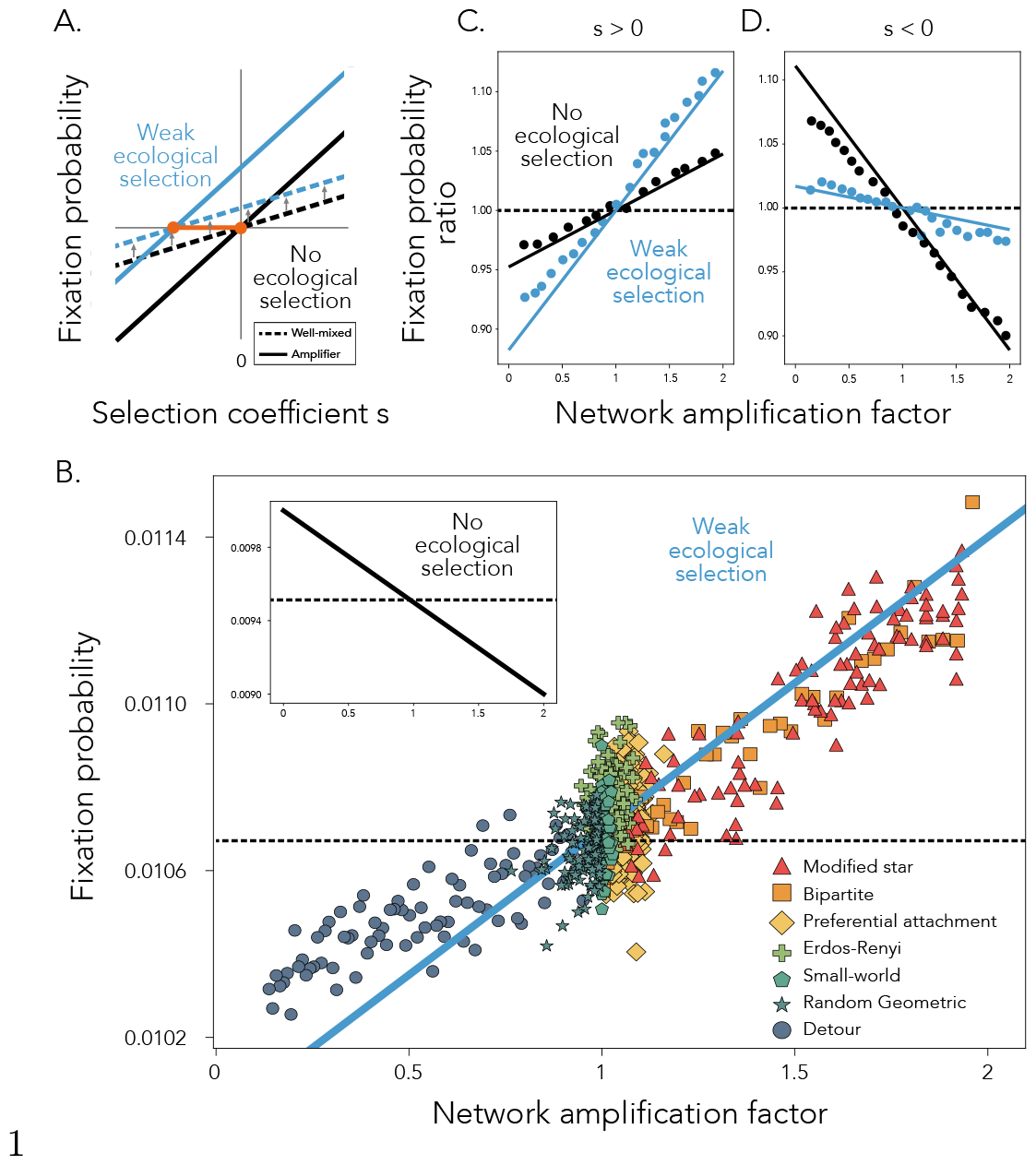
The role of network topology in shaping mutant fixation probabilities for generalist populations. **Panel A** shows an illustration of the effect of weak ecological selection on the probability of fixation. Fixation probability is shown on the y-axis and selection strength on the x-axis. One set of black dashed (well-mixed) and continuous (amplifier) lines show the fixation probability under no ecological selection, while the other set in blue showcases the regime with weak ecological selection. The orange points show the points of intersection for the two ecological regimes. **Panel B** Fixation probability on the y-axis as a function of network amplification factor on the x-axis, across network families. Each dot represents the fixation probability of a single network, calculated using 10^6^ simulation runs. The dashed line represents equation (5) for a well-mixed population, while the solid blue line shows equation (6) for graph topologies. Here, *s* = − 0.001, *N* = 100 and *α* = 0.53 (blue solid line) and *α* = 0.5 for the insert. In **Panels C** and **D**, dots show the ratio of fixation probabilities across network families and the well-mixed population baseline, as a function of network amplification factor, for a beneficial (*s* = − 0.001, **Panel C**) and deleterious (*s* = 0.002, **Panel D**) mutant, estimated using 10^6^ simulation runs. Lines show the analytic approximations using equations (5) and (6). Here, *N* = 100 and *α* = 0.525 (blue lines).

In what follows, we analyze how the network structure changes the mutant’s fixation probability and fixation time, compared to a well-mixed population, as it now operates on two fitness-affecting axes: the constant pure selection coefficient, *s* and the ecological niche overlap, *α*, which gives rise to frequency-dependent selection. We start by describing the eco-evolutionary dynamics in the limit of weak ecological selection, with generalist ecotypes that utilize both resources almost equally and show that the mutant dynamics on the network are shaped by a combined effective selection coefficient that captures the interplay of the two fitness coefficients. We then show that, at the opposite limit of strong ecological selection, with specialist species that have little niche overlap, mutant spread and fixation are shaped by a tradeoff between establishment and conditional fixation probabilities, with the two fitness coefficients playing sequential roles. We then discuss how these two edge regimes provide intuition on the shape of the dynamics observed for ecological interactions of any strength and show that the addition of global frequency-dependent ecological selection pressure can reverse the roles of network topologies traditionally assumed to amplify (and conversely, the ones assumed to suppress) selection.

### The regime of weak ecological selection

In the regime of weak ecological selection, the variants are generalist species that rely on both available resources almost equally. At a big picture level, we show that the interplay between the pure fitness selection coefficient *s* and the ecological selection *α* gives rise to a combined effective selective coefficient for the mutant, which we write out below. The network topology further amplifies or suppresses this translated effective selection coefficient (**Figure 1A**). This can give rise to parameter regimes where the ecological selection makes ‘amplifier’ network topologies amplify deleterious mutants (*s <* 0) and ‘suppressor’ topologies suppress deleterious mutants, relative to a well-mixed population, in contrast to results observed without ecological selection (*α* = 0.5).

In this regime, the global frequency-dependent ecological component of mutant fitness pushes the mutant frequency towards an internal equilibrium, while the constant pure fitness component biases this equilibrium in favor of the fitter ecotype. The mutant frequency at equilibrium is given by

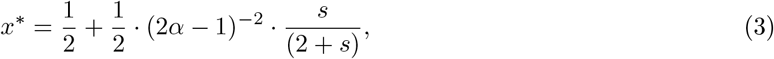

for *α >* 0.5 (derivation in **Appendix 1**). When *α* = 0.5, there is no equilibrium because there is no ecological selection. When *s* = 0, the equilibrium frequency is 0.5 for all *α* because neither variant has a fitness advantage. The mutant is more abundant at equilibrium (*x*^*∗*^ *>* 0.5) if *s >* 0 and the wild-type is more abundant (*x*^*∗*^ *<* 0.5) if *s <* 0.

In the limit of weak ecological and small pure fitness *s*, strong genetic drift makes this equilibrium unstable and the dynamics appear neutral-like. The interplay between the pure fitness coefficient and ecological selection gives rise to a combined effective selective coefficient that is approximately

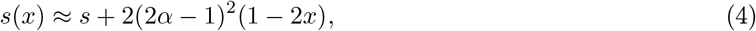

where x is the mutant frequency in the population (see **Appendix 2** for this derivation). Therefore, in a well-mixed population, the fixation probability of the mutant becomes approximately

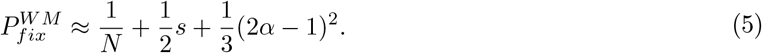

The original population under frequency-dependent selection behaves effectively as one under constant selection, with 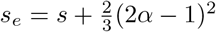.

Since the network structure reshapes the mutant’s selection coefficient by increasing or diminishing it proportional to the amplification parameter of the graph, *a*_*Bd*_, the mutant’s new effective selection coefficient becomes *a*_*Bd*_*s*_*e*_. For a spatial structure G, the fixation probability of the new mutant can therefore be approximated as

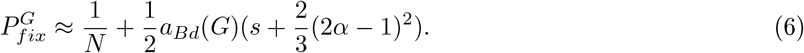

**Figure 1B** shows this analytic approximation of the fixation probability (blue line) as a function of *a*_*Bd*_(*G*) on the x-axis, together with simulation results, for a variety of graph families, both suppressors and amplifiers of selection. As a comparison, the insert shows results for the same deleterious mutant with *s* = −0.001, in the absence of ecological selection (*α* = 0.5). When *s*_*e*_ is net positive despite a negative pure *s*, i.e. the beneficial contribution from ecological selection pressure is greater than the deleterious contribution from constant selection *s*, an amplifier will promote this deleterious mutant, compared to a well-mixed population. Concretely, this happens when 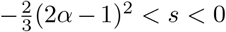. Outside of this region, the effect of the graph topology does not change with the addition of eco-evolutionary interactions, however the strength of amplification and suppression does change. For beneficial mutants, ecological selection increases the strength of both amplification and suppression (**Figure 1C**), while the opposite is true for deleterious mutants (**Figure 1D**).

Note that the effective selection experienced by the mutant is equal to the selection it experiences when it has frequency 1*/*3 in the population, i.e. 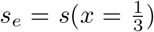, as previously observed in other models (Nowak et al., 2004; Ohtsuki et al., 2007a). We can utilize this result to further generalize the weak selection approximation to cases with unequal resource consumption rates and/or asymmetric resource supply rates (**Supplementary Material Figure S4**).

From this result, it also follows that graphs where each node has the same number of neighbors (lattice structures, or more generally, k-regular graphs) obey the isothermal property, since their amplification factor is equal to 1, and they do not change probabilities of fixation compared to a well-mixed topology (**Supplementary Figure S1**). The isothermal theorem does not usually hold for k-regular graphs in prior models of game-theoretic frequency-dependent dynamics (Ohtsuki et al., 2006). This is due to the fact that in these prior models frequency dependence is local and the fitness of an individual depends on the frequency of its own type amongst its neighbors (Allen et al., 2017).

### The regime of strong ecological selection

In the edge regime of strong ecological selection both variants are specialist species that consume different resources, with only some niche overlap. The strong frequency-dependent ecological selection makes the mutant persist at the equilibrium frequency for much longer, before eventual fixation or loss from the population (**Figure 2A**). In what follows, we discuss how network topology, in interplay with the strength of niche differentiation, shapes the time the population spends at this interior equilibrium and how that affects fixation probabilities in this regime. It is convenient to study the fixation process as the outcome of two independent processes: the probability of the initial establishment of the mutant in the population, followed by the probability of conditional fixation, conditional on mutant frequency starting from the equilibrium frequency (Grimm and Wissel, 2004). We define establishment as the change from one mutant to *Nx*^*∗*^ mutants (internal equilibrium) and conditional fixation as the change in mutant frequency from *Nx*^*∗*^ mutants to N mutants (fixation) (**Figure 2A**).

**Figure 2.**
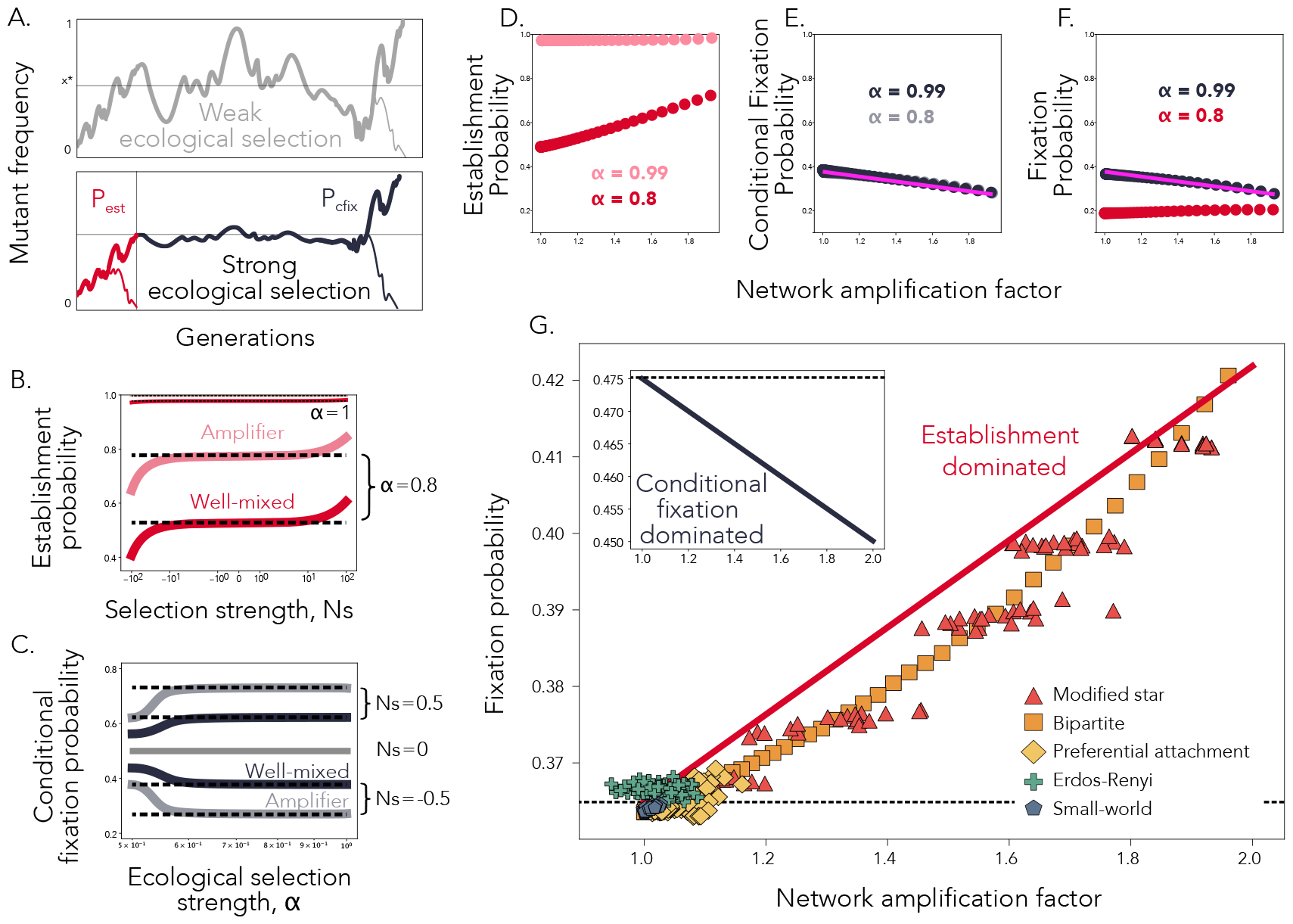
The role of spatial structure in specialist populations. **Panel A** shows toy representations of the mutant trajectory in the generalist (grey) and in the specialist (red/black) regimes. **Panel B** shows the theoretically exact establishment probabilities for well-mixed (solid lines, darker red) and an amplifier (solid lines, lighter red) population (equations (A30) and (A35)). Here, *a*_*Bd*_ = 2, *N* = 500 and *α* = 0.8. The slight change in the establishment probability is also showcased for *α* = 0.99. The corresponding analytic approximations (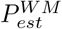 and 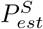) are shown using black dotted lines. **Panel C** shows the theoretically exact conditional fixation probabilities (equations (A29) and (A34)) for a well-mixed (solid lines, darker color) and an amplifier (solid lines, lighter color) population, with amplification factor 2. Here, *N* = 500 and three different values of *Ns*, as depicted in the figure. The corresponding analytic approximations (equations (11) and (12)) are shown using black dotted lines. **Panel D** shows establishment probabilities, **Panel E** shows conditional fixation probabilities, and **Panel F** shows total fixation probabilities, as a function of amplification factor, for various PA star networks, computed numerically for two different values of *α*, as shown, and *s* = − 0.01. The solid pink line shows the analytic approximation for the conditional fixation probability given by equation (12). **Panel G** Dots show an approximation of fixation probability using the approach described in the main text on a diverse array of amplifier topologies with varying amplification factors. Here, *N* = 100, *s* = − 0.001 and *α* = 0.9. The solid red line shows equation (13) and the dotted line showcases the well-mixed result as comparison. The insert shows the analytic approximation of the fixation probability, given by equation (12), for *s* = − 0.001 and *α* = 1 (solid black line). The dotted line again shows the well-mixed fixation probability.

In a well-mixed population, the fixation probability is the product of the probabilities of success of both processes: *P*_*fix*_ = *P*_*est*_*P*_*cfix*_. In spatially-structured populations, however, we have to account for all the different ways to arrange *Nx*^*∗*^ mutants on the graph, at equilibrium. Let **A** be a N x N^*^ matrix in which element (*i, j*) is the probability of establishing in the *j* graph equilibrium configuration, given a mutant initialized in node *i*. Here, N^*^ is the total number of spatial equilibrium configurations, 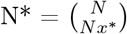. Let **B** be the vector of size N^*^, with element *i* equal to the probability of fixation, conditional on establishment in the *i* equilibrium configuration. The fixation probability vector, where element *i* is the fixation probability given mutant initialization in node *i*, is then given by **A** · **B**. The establishment probability for a graph *G*, i. e. the probability of the mutant establishing in any of the N^*^ equilibrium configurations given that it is initialized on a randomly selected node, becomes

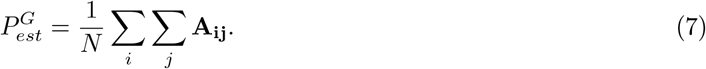

Similarly, the conditional fixation probability on a graph can be written as

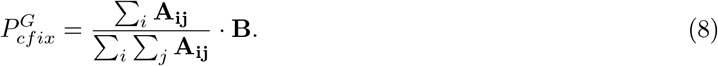

The 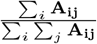 term is a N^*^-element vector in which the *i*-th element is the probability of establishing in equilibrium configuration *i*, given that establishment is successful and is a weighted average of the conditional fixation probabilities corresponding to each equilibrium state, where the weights are the relative probabilities of establishment in each of those states. Therefore, for a graph *G*, similar to the well-mixed case, we find that the mutant probability of fixation is the product of the probabilities of success of both establishment and conditional fixation,

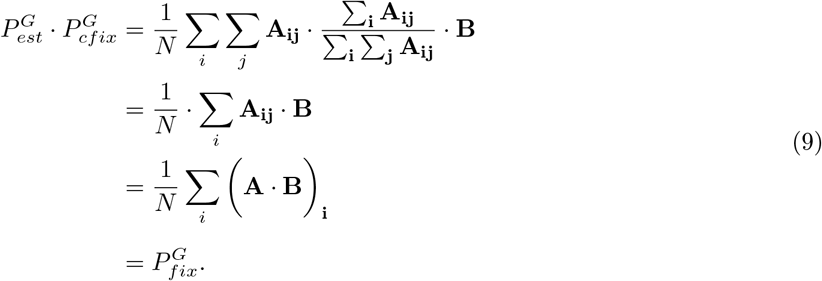

It is therefore sufficient to study these two probabilities independently. In what follows, we discuss how to derive intuitive approximations and also exactly compute these probabilities using exact equations for special case spatial structures (see more in **Appendix 4**), and methods for probabilities of Markov Chain absorption for a broader class of spatial structures (Tolver, 2016; Hindersin et al., 2016).

#### The probability of establishment

For a well-mixed topology, in the limit of weak selection *s* and large population size *N*, the establishment probability can be approximated as

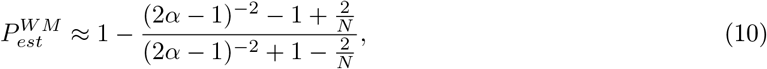

(see derivation in **Appendix 5**). This approximation is useful, because it shows that *P*_*est*_ does not depend on the pure fitness coefficient, *s* (**Figure 2B**). Alternatively, we can write 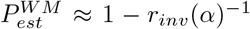, where *r*_*inv*_(*α*) is the ecological fitness of the mutant when it has frequency 1/N in the population and note that 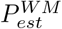 is equivalent to the fixation probability of a mutant with relative fitness *r*_*inv*_(*α*). This makes intuitive sense, since *r*_*inv*_(*α*) is the approximate fitness of the mutant in the initial stochastic regime, where genetic drift dominates and before the invasion fitness is significantly impacted by the frequency of the mutant in the population (Good et al., 2018). As in the weak ecological selection regime, establishment can be associated with an effective constant fitness.

A network population structure reshapes this effective fitness *r*_*inv*_(*α*). However, in contrast to the previous regime where the ecological selection pressure is weak, in this case it is harder to obtain general results, except for highly-symmetric topologies. For example, star-structured amplifiers reshape fitness *r* → *r*^2^ (Lieberman et al., 2005) and we can therefore approximate the establishment probability on the star as 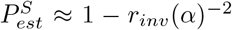 (see also **Figure S2**). Note that as *α* approaches 1, ecological selection becomes so strong that establishment occurs almost deterministically, and therefore *P*_*est*_ saturates at 1 for both the well-mixed and star-structured populations (**Figure 2B**).

#### The probability of conditional fixation

The conditional fixation probability is, in contrast, dominated by the pure fitness coefficient *s* and we can write

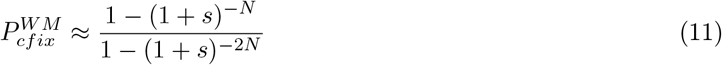

(refer to **Appendix 6** for this derivation). During conditional fixation, spatial structure therefore acts on the pure selection coefficient *s* (**Figure 2C**) and, for a star-structured population, we can write in closed form 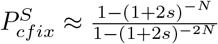 (see also **Figure S2**). Note that the star topology boosts the conditional fixation of beneficial mutants and suppresses the fixation of deleterious mutants, as in the constant selection case.

Because conditional fixation is independent of the ecological selection coefficient *α*, more general derivations are also possible because we can use the network amplification parameter *a*_*Bd*_ as it reshapes only the constant fitness component *s*. For a graph *G* with amplification factor *a*_*Bd*_(*G*), the probability of conditional fixation can be written as

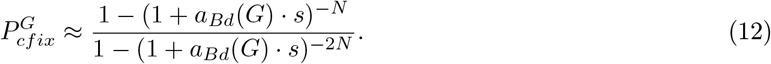

#### The total probability of mutant fixation

While the frequency dependent dynamics makes the probability of establishment more difficult to analytically compute for topological structures of arbitrary complexity, similar to the case of local frequency dependent dynamics (Hadjichrysanthou et al., 2011; Altrock et al., 2017), in what follows we show how we can use a combination of simulations and analytic approximations to calculate mutant fixation probability for arbitrarily heterogenous topologies.

For models without global frequency dependence, the probabilities of mutant fixation linearly increase (for *s >* 0) and decrease (for *s <* 0) as a function of the network amplification factor (see insert in **Figure 1B** for an example *s <* 0 case). In our model, this result holds for advantageous mutants with *s >* 0 because the effect of the topology is the same for both establishment and conditional fixation, i.e. if a spatial structure increases the establishment probability of the mutant, it will also increase its conditional fixation probability, and vice versa. However, here we show that this result does not hold for weakly deleterious mutants with *s <* 0 and the relationship between amplification and mutant fixation probability is fundamentally determined and reshaped by the strength of ecological selection *α*.

We first study a family of graphs, preferential attachment (PA)-star graphs, for which establishment, conditional fixation, and fixation probabilities can be computed exactly using numerical methods, due to their degree of symmetry (Hindersin et al., 2016; tka) (see **Materials and Methods**). Even for the case of a deleterious mutant *s <* 0, the probability of establishment increases with the network amplification factor because the effective constant fitness associated with establishment is beneficial (*r*_*inv*_(*α*) *>* 1) (**Figure 2D**). As the population becomes increasingly specialist (*α* → 1), the establishment probability saturates at 1 for all amplification factors, i.e. spatial structure has a negligible effect on establishment in this regime.

Separately, the conditional fixation probability decreases with the amplification factor because the effective constant selection coefficient associated with conditional fixation (*a*_*Bd*_*s*) is negative (**Figure 2E**).

Therefore, the overall fixation probability increases with network amplification factor only in an “establishment dominated regime” (red line, *α* = 0.8 in **Figure 2F**), where the amplification property of the graph topology increases establishment by a larger amount than it decreases conditional fixation. In contrast, in a “conditional fixation dominated regime”, amplification has almost no effect on establishment, but still decreases conditional fixation by the same amount. Therefore, in this regime, mutant fixation probability is approximately equivalent to the conditional fixation probability, 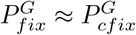 and it is decreased for networks that are traditionally classified as ‘strong’ amplifiers (black line with simulation results and pink numerical approximation results for *α* = 0.99 in **Figure 2F**).

We can analyze the probability of fixation for a broader class of topologies by computing the establishment and conditional fixation through a combination of simulations and analytics: since the mutant is under strong ecological selection until the equilibrium is reached, the establishment is fast and can be estimated through simulations for more heterogeneous topologies. Separately, equation (12) provides an approximation for the conditional fixation. In **Figure 2G** we show that, in the establishment-dominated regime (*α* = 0.9), the fixation probability of the mutant increases with the network amplification factor across a wide array of network families. In contrast, in the regime where the probability of conditional fixation dominates (*α* = 1), network amplification factor decreases the fixation probability (inset **Figure 2G** for the same selection coefficient *s* = −0.001).

In this case, the relationship between the fixation probability and the amplification factor of the network is approximately linear. Since the exact fixation probability can be computed analytically for the well-mixed and star populations, we can use these two points to write a linear approximation for the fixation probability for a spatial structure of arbitrary amplification factor (red line in **Figure 2G**)

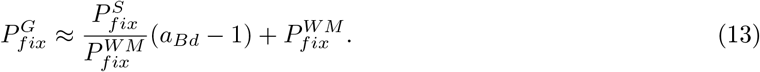

Because fixation is associated with two effective selection coefficients, one for establishment and one for conditional fixation, there also exist parameter regimes where the larger the probability of establishment, the smaller the probability of conditional fixation, and vice versa (**Figure 3A**), leading to a establishment-conditional fixation tradeoff. A consequence of this tradeoff is that, for weakly deleterious mutants, a nonlinear relationship between the amplification factor of the network and the mutant fixation probability can emerge and fixation probabilities can be maximized by spatial structures with intermediate amplification factors (**Figure 3B**).

**Figure 3.**
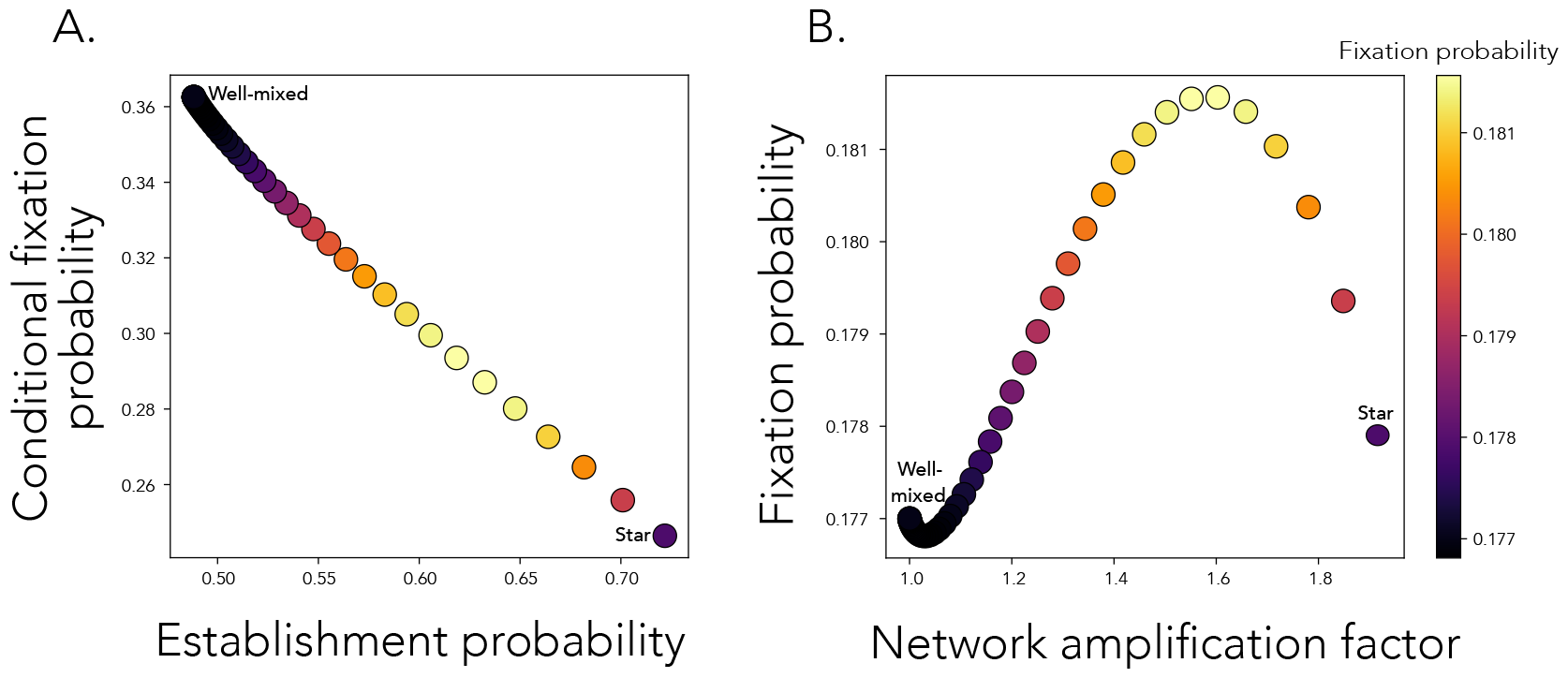
There exist parameter regimes with establishment - conditional fixation tradeoffs. **Panel A**: Each point shows the conditional fixation probability as a function of the establishment probability for amplifier networks with amplification factor as on the x-axis in Panel B. The total fixation probability is shown by the color of the point, as in the colorbar. In **Panel B**, each point shows the total fixation probability as a function of the amplification factor of the network. For both panels, *α* = 0.8 and *s* = −0.012.

### The strength of resource niche competition determines whether a spatial topology amplifies or suppresses weakly deleterious mutants

A key finding resulting from the analyses above is that topologies that are known amplifiers of selection can also amplify deleterious mutants (*s <* 0) and the strength of ecological selection determines the range of parameters where this result holds (**Figure 4A**). This unintuitive, nonlinear behavior is in stark contrast to the behavior observed under constant selection models, where amplifier networks decrease the fixation probability of deleterious mutants.

**Figure 4.**
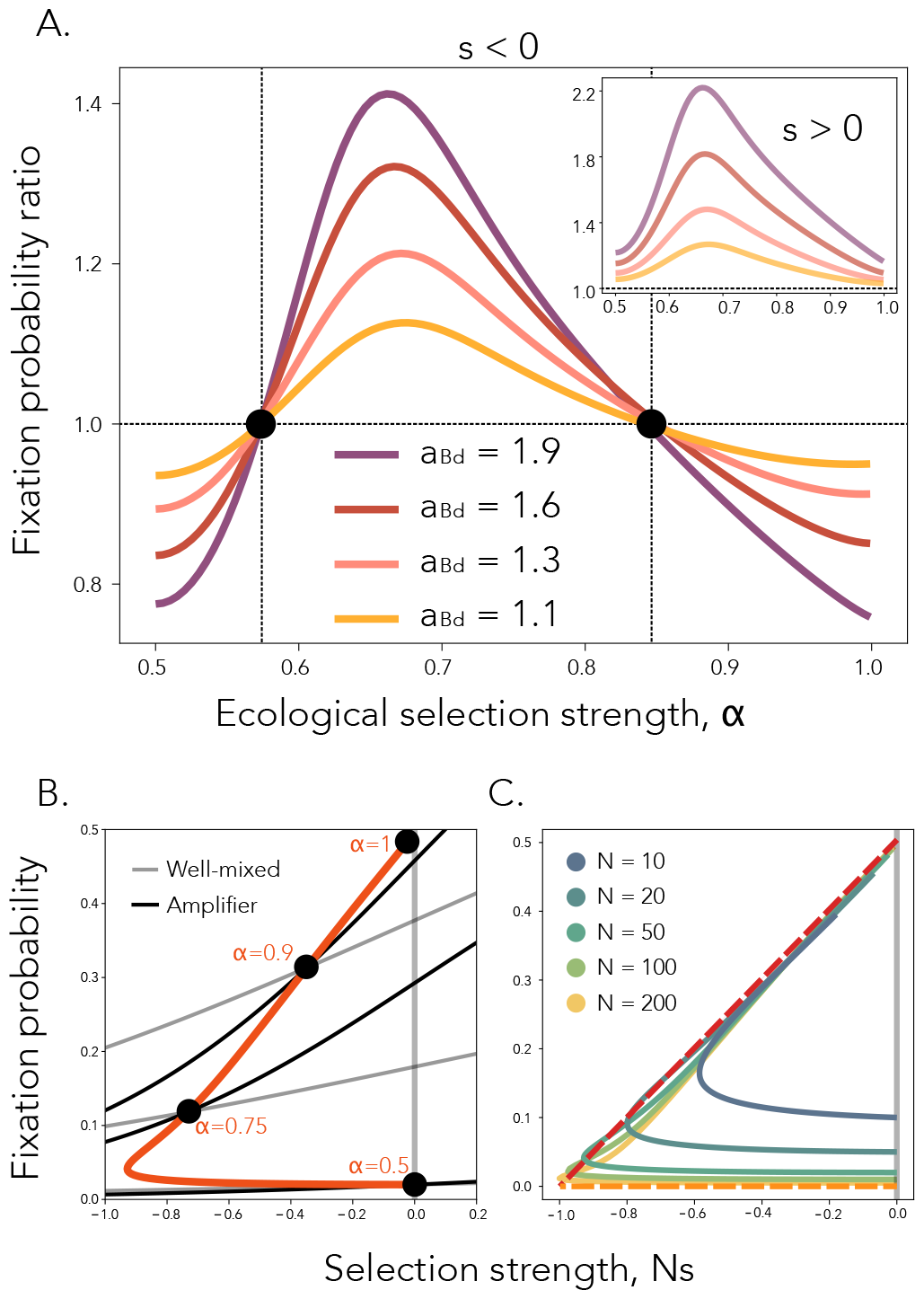
Ecological interactions can reverse the role of amplifiers and suppressors for weakly deleterious mutants. **Panel A**. Ratio of fixation probabilities between various amplifier networks and a well-mixed population. Here, *N* = 50 and *s* = − 0.01. The insert shows a beneficial mutant with *s* = 0.01. **Panel B**. Comparison of probabilities of fixation as a function of mutant selection strength for a well-mixed population (grey lines) and an amplifier of selection (black lines, amplification factor equal to 1.92), for four different values of ecological strength *α*, as written in the figure. The orange line follows the exact point of intersection between the well-mixed and amplifier populations, as ecological strength *α* is continuously varied. For *α* = 0.5, we recapture previous results from evolutionary graph theory, where the intersection point is at *Ns* = 0. Here, fixation probabilities are computed exactly, as described in **Appendix 4**. Population size *N* = 50. **Panel C**. The points of intersection of the fixation probability between well mixed and a star population, for varying population sizes *N*, as depicted in the legend. The orange dotted line shows the approximation for 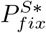 in the generalist regime (**Appendix 3**) and the red dotted line shows the approximation for 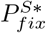 in the specialist regime (see **Appendix 7**).

**Figure 5.**
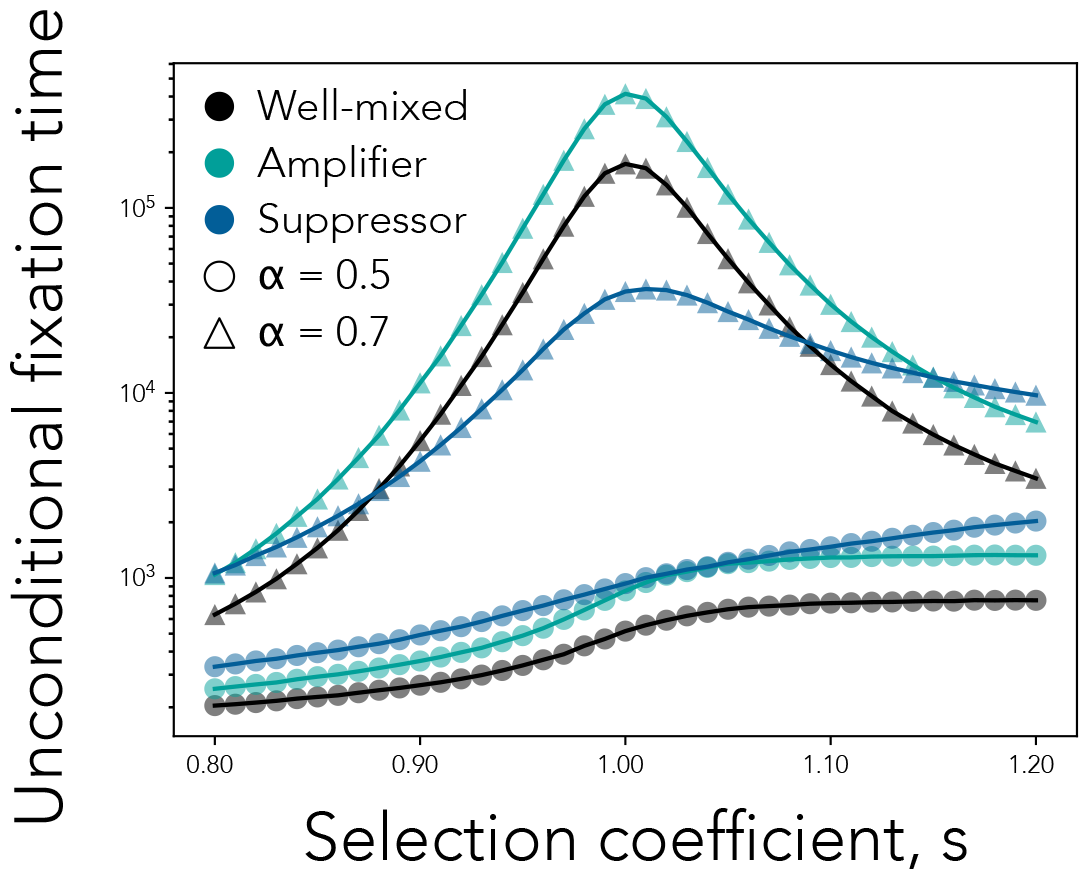
Well-mixed populations do not minimize fixation time. We plot the unconditional fixation time as a function of the pure selection coefficient *s* for a well-mixed population, a suppressor, and an amplifier network. Here, *N* = 100. Each point of the graph shows the unconditional fixation time for a single network for a specific values of *s* and *α*, using 10^6^ simulation runs. Solid lines connect points corresponding to the same network. The shape of the point represents the value of ecological selection and the color represents the network type, as shown in the legend. In contrast to dynamics observed in the absence of ecological selection (*α* = 0.5), for *α >* 0.5, there exist network topologies that can decrease time at equilibrium and time to fixation, compared to well-mixed populations.

Starting in the limit of *α* = 1, when the two ecotypes have no niche overlap, the initially rare mutant will experience strong positive ecological selection, because it consumes a resource that is completely unused by the wild-type population. Therefore, establishment occurs almost deterministically, regardless of the spatial structure of the population, i.e. 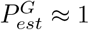, and the overall probability of fixation is determined by the conditional probability of fixation, 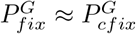 and this represents a conditional fixation dominated regime.

As *α* decreases, establishment is no longer deterministic and spatial structure will start contributing to the probability of establishment. If the amount by which the graph topology boosts mutant establishment outweighs the amount by which it suppresses conditional fixation, we enter the establishment dominated regime, and the mutant fixation probability on the graph becomes larger than that on a well-mixed structure (second crossing point in **Figure 4A**). As *α* keeps decreasing, we enter the generalist regime in which, for strong enough ecological differences, the effective selection coefficient 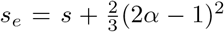 is still positive, so the amplifier topology keeps amplifying the deleterious mutant. In the limit of no ecological selection, as *α* approaches 0.5, the effective selection coefficient once more becomes net deleterious and the spatial structure again suppresses the new mutant in the population (first crossing point in **Figure 4A**). In contrast, amplifiers always increase the fixation probability of beneficial ecotype mutants, in line with previous theory under constant selection (*s >* 0 insert **Figure 4A**).

Another way to understand this result is through the lens of how *α* determines the intersection point between the fixation probability on a graph topology and on a well-mixed topology as a function of the strength of selection *Ns*. Evolutionary graph theory with constant selection defines amplifiers as topologies that increase the fixation probability of beneficial mutants and decrease that of deleterious ones. Therefore the respective probabilities of fixation intersect at *Ns* = 0 (see **Figure1A** black lines). With the addition of global frequency-dependent ecological interactions, this intersection point is shifted to negative values of *Ns*. Specifically, there exists a region of selection against weakly deleterious mutants where ecological selection reverses the evolutionary effect of the spatial structure and amplifiers can favor weakly deleterious mutations and boost their spread compared to well-mixed populations. The extent of this region is maximized for an intermediate value of *α*, such that amplifiers disfavor all deleterious mutations either in the absence of ecological differences, for *α* = 0.5, or for complete lack of niche overlap and very strong ecological selection, *α* = 1 (**Figure 4B**). We find that strong amplifiers like the star graph can increase the fixation probability, relative to well-mixed, of deleterious mutants with up to *Ns* ≈ −1 even for small population sizes (**Figure 4C**). We also show results for suppressor topologies in **Supplementary Figure S3**.

### The time to fixation

As the strength of ecological selection increases, the time to fixation of the new mutant also increases across all spatial structures. This is because the frequency-dependent selection increases the pull towards the unstable interior equilibrium. In the absence of ecological selection, it is known that the well-mixed populations minimize fixation time of new mutants in the population (tka). Here however, we observe that, with ecological selection, it is possible for a population on a suppressor network to have a significantly shorter fixation time than the well-mixed population. There are two reasons for this. Firstly, when a mutant first appears in small quantities, in most cases it will immediately drift to extinction. However, the probability of immediate mutant loss is even larger on suppressor networks compared to well-mixed populations because suppressors decrease effective selection. Secondly, if the mutant successfully reaches equilibrium, the amount of ecological selection determines the strength of negative frequency-dependent selection pushing the ecotype frequencies towards the ecological equilibrium. Since suppressors decrease effective selection, the equilibrium is intrinsically less stable in these populations compared to well-mixed populations and the force of drift can more easily push the mutant towards the two absorbing equilibria.

## Discussion

We introduce a mathematical framework that combines evolutionary graph theory with a resource competition model to study how heterogeneous topological structure shapes evolutionary dynamics under global frequency-dependent ecological interactions. We analyze the probability of fixation of a new ecotype mutant, characterized by two fitness axes: a pure fitness axis and an ecological fitness axis, driven by the extent of niche separation. This model can be equivalently expressed as an evolutionary game on graphs in which payoff matrix elements are functions of the two parameters that control the strengths of ecological and pure selection (Nowak, 2006; Allen et al., 2017). A main difference between our model and previous literature on local frequency dependence dynamics on graphs is that here all resources are supplied by the environment and are not a diffusible public good produced through cooperation dynamics. Although some limited work exists in cases where replacement and frequency-dependent interactions occur over the entire network, both replacement and interaction networks are assumed to be sparse (low mean degree) (Ohtsuki et al., 2007b). This global frequency-dependence assumption also differentiates our model from others that assume each spatial niche hosts only one type of resource (Kaveh et al., 2020; Nemati et al., 2023).

This work builds on prior models of ecotype dynamics, each separately incorporating (and potentially examining in more detail) some of the integral parts of the model we build here. In some prior studies, all ecotype variants are assumed to be evolutionarily neutral (Posfai et al., 2017; D’Andrea et al., 2020; Weiner et al., 2019) and/or analyze ecotype dynamics only in populations with highly symmetric, lattice-type spatial structures (Weiner et al., 2019). Separately, other previous models of consumer-resource dynamics on more complex graph structures assume a single predefined, spatially-heterogenous resource that all variants rely on, which effectively means they all occupy the same ecological niche (Kaveh et al., 2020; Nemati et al., 2023). Other models allow for intrinsic fitness differences between different ecotype strains, however do not account for spatial structure (Good et al., 2018).

While in most prior models that incorporate ecological niche differences, the focus is on understanding the population at equilibrium and factors such as the richness (Huang et al., 2012) and abundance distributions of species (Phillips et al., 2006), an important distinction to our analysis is that we do not assume the limit of infinite population size and, in our model, genetic drift will eventually push the mutant to either fixation or loss from the population. This is similar to Good et al. (2018), where ecotype fixation is also possible and may occur if a sufficient number of pure fitness-modifying mutations accumulate within a clade. Prior work has also only considered how network structure shapes either 1) constant selection dynamics in isolation or 2) frequency-dependent dynamics, without recognizing that even variants under frequency-dependent selection can carry an intrinsic constant fitness axis that spatial structure can act on.

Using the case without ecological differences as an explicit limit of our model, we show that the effects of an ecological, frequency-dependent fitness axis can affect the evolutionary properties of the network in non-intuitive ways. While amplifier spatial networks are known to suppress deleterious mutants (Lieberman et al., 2005), here we show that weakly deleterious mutants, which can exploit ecological niche differences, can have increased fixation probabilities on amplifier networks, compared to well-mixed populations. We also show that the reverse is true for suppressor networks of transmission - the coupling of spatial topology and ecological selection can suppress weakly deleterious mutants. We show that the extent of this reversal in spatial structure effect is a non-monotonic function of niche overlap (i.e. strength of ecological selection) and we explain this result by considering the edge cases of weak and strong niche overlap.

In the regime of generalist populations, we can write out the effective fitness coefficient of the mutant as a function of the initial pure selective coefficient, the ecological diversification strategy and the evolutionary topological properties of the network. The alternative edge case of little niche overlap (specialist ecotypes) corresponds to an evolutionary game on a graph under strong selection, which is generally not well understood. The fixation probability under arbitrary selection intensity is only known for some symmetric spatial structures (Ohtsuki and Nowak, 2006; Hadjichrysanthou et al., 2011; Broom and Rychtář, 2008). Here we show that the net fixation probability can be decomposed into establishment and conditional fixation probabilities and that 1) establishment depends only on the strength of the frequency-dependent (ecological) selection and 2) conditional fixation depends only on the strength of the frequency-independent (pure) selection. An analagous result is known for the fixation time under weak selection: 1) the conditional fixation time depends only on the frequency-dependent selection and 2) the unconditional fixation time depends only on frequency-independent selection (Altrock and Traulsen, 2009). Distinct establishment and conditional fixation subprocesses in specialist populations cause significant departures from evolutionary dynamics on graphs under effectively constant selection. For example, while the relationship between amplification and fixation probability is expected to be monotonic (Kuo et al., 2021), we find that, for certain parameter regimes, it is graphs with intermediate amplification factor that can maximize the overall fixation probability of weakly deleterious mutants in specialist populations, because they achieve the best tradeoff between establishment and conditional fixation.

For analytical conveniency and to curtail the number of parameters, our work has focused on a highly simplified model, which omits many of the complicating factors expected in either natural or laboratory settings. The optimal model choice depends on which parameters can be measured and future work could explore the effects of spatially-heterogenous resources (Nemati et al., 2023; Fu et al., 2015), the effects of cross-feeding (Archetti and Pienta, 2019; D’Souza et al., 2018), higher-order multi-species interactions (Bairey et al., 2016) and other more diverse modes of ecological interaction (Luhring and DeLong, 2020). Our model also assumes that pure fitness differences are directly linked to diversification-related mutations, rather than arising independently through secondary mutations. In addition, the strength of ecological selection is not allowed to evolve over time. These are all assumptions that need to be relaxed for a systematic understanding of eco-evolutionary dynamics. Our results provide an initial general framework for integrating ecological differences, population-genetic processes and complex patterns of transmission to study spatially heterogenous evolving ecosystems and lay the groundwork for engineering structures that select for communities of ecological, medical, or industrial utility.

## Materials and Methods

Understanding which network properties shape eco-evolutionary dynamics is complicated by the fact that these properties are often correlated, hard to tune independently and differ across many network families. In previous work, we showed that linking network topology to evolutionary dynamics can be understood and analytically quantified by computing the network amplification factor, without having to keep track of all the lower-levels network properties (Kuo et al., 2021; Kuo and Carja, 2024).

Here we use the approximation for the amplification factor from Kuo et al. (2021). We can also compute the amplification factor empirically: using the Birth-death process, and starting with one single mutant with fitness (1 + *s*) invading a population of wild-type individuals with fitness 1, the amplification factor can be computed using the definition from (Lieberman et al., 2005) and solving

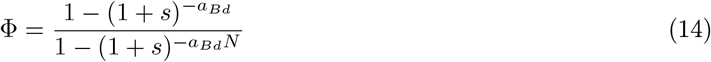

for *a*_*Bd*_, where Φ is the mutant’s fixation probability.

In this study, we use a combination of well-known complex network families using built-in generators from NetworkX (Hagberg et al., 2008) and also design graphs that allow us to tune the amplification factor and enable numerical tractability for our model, as detailed below.

### Preferential attachment graphs

For graphs with preferential attachment, nodes are added sequentially starting from a set *m* of initial nodes, until the population reaches size *N*. Each new node is added to the network and connected to other individuals with a probability proportional to the individual’s current degree to the power of a given parameter of preferential attachment, *β*. We use *m* ∈ {3, 5, 20}, and *β* ∈ [−3, 3].

### Preferential attachment stars

We introduce a family of symmetric preferential attachment-type graphs which allow exact computation of the mutant fixation probability, using numerical methods for absorbing Markov chains (Hindersin et al., 2016; Tolver, 2016). These graphs strike a balance between exhibiting enough spatial heterogeneity, but also enough symmetry for analytical convenience. Specifically, a *N* -node PA star contains: 1) *i* core nodes, each connected to every other node in the graph, and 2) *N* − *i* leaf nodes, connected to all *i* core nodes. When *i* = 1, we recover the star graph, and when *i* = *N* − 1 we recover a well-mixed graph. Because only two variables are needed to represent any configuration of mutants on this type of graph, the mutant frequencies on core and leaf nodes, the symmetric structure of a PA star reduces the state space of the Markov process from 2^*N*^ states to (*i* + 1) · (*N* − *i* + 1) states, making numerical calculation possible. A PA star with N nodes and i core nodes is equivalent to a preferential attachment network with *m* = *i* and *β* = ∞.

### *k*-regular graphs

In a k-regular graph, every node has the same number of neighbors, *k*. We generate *k*-regular graphs using built-in generators from NetworkX, for different values of *k*, as presented in the text.

### Erdős Rényi random networks

The Erdős Rényi network is initialized with N disconnected nodes. Every pair of nodes is then connected independently with a probability *p*. We use NetworkX to generate these networks.

### Small world networks

The small world network is initialized with a regular ring lattice network of degree *k*. For every node, each of its neighbors are rewired with probability *p*. We use *k* ∈ {8, 12, 16}, and *p* ∈ [0, 1]. We generate small world networks using built-in generators from NetworkX.

### Bipartite graphs

Bipartite graphs have two sets of nodes *n*_1_ and *n*_2_ such that edges connect nodes across the two sets. We vary *n*_1_ from 1 to 50, and *n*_2_ = 100 − *n*_1_. We generate small world networks using built-in generators from NetworkX.

### Random geometric graphs

Random geometric networks are spatial networks with nodes sampled in a 2-dimensional space; we sample x coordinates for 50 nodes from 𝒩 (0, 0.25) and another 50 from 𝒩 (3, 0.25) and y-coordinates for all 100 nodes from 𝒩 (0, 0.25). We start with the lowest cut-off radius, resulting in a connected graph, and increase the cut-off radius until the complete graph is formed.

### Detour graphs

Detour graphs are initialized with a complete graph of size *n*_1_. Then one edge is replaced with a line graph of length *n*_2_. We use *n*_1_ ∈ [3, 99], and *n*_2_ = 100 − *n*_1_.

### Modified star graphs

The star graph consists of one center node connected to *N* − 1 outer nodes. We generate star graphs using built-in generators from NetworkX. Modified star graphs are initialized with a star graph of size N. Then *m* random edges are added to the graph. We use *m* = [50, 100, 200, 300].

## Acknowledgments

This research was done using resources provided by the Open Science Grid, which is supported by the National Science Foundation award 1148698, and the U.S. Department of Energy’s Office of Science.

## Funding

We gratefully acknowledge support from the NIH National Institute of General Medical Sciences (award no. R35GM147445), the NIH T32 training grant (no. T32 EB009403) and the United States-Israel Binational Science Foundation (award no. 2019266).

## Appendices

### A1: Deriving the mutant frequency at equilibrium

From equation (1), the steady state concentrations of the two resources are

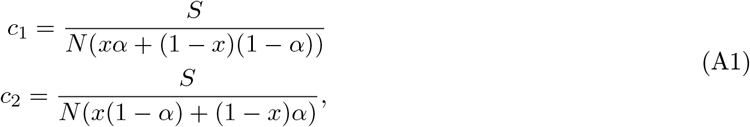

where *N* is the population size and *x* is the mutant frequency in the population. Therefore, we can write the ecological mutant fitness as

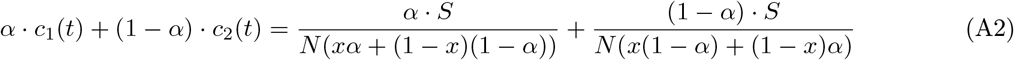

and the wild-type fitness as

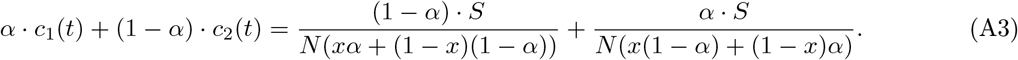

Therefore, the relative ecological fitness of the mutant is given by

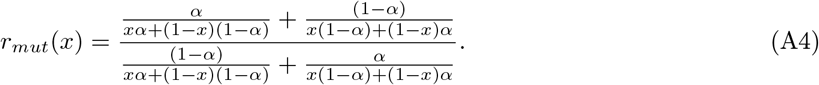

To simplify the analysis, we show that relative mutant fitness can be written in a hyperbolic form as

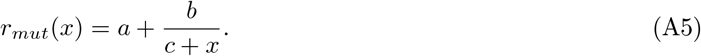

To determine the coefficients in (A5), we use the following three conditions that *r*_*mut*_(*x*) must satisfy:

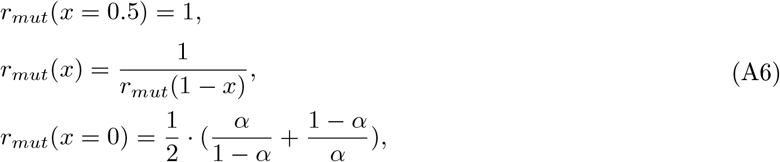

where the last condition follows from (A4). We can write

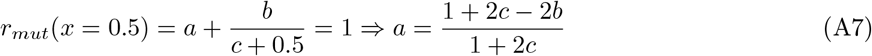

and plugging back into (A5),

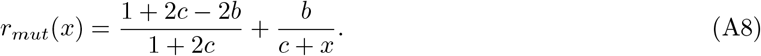

From the middle condition in (A6), for *x* = 1, we get

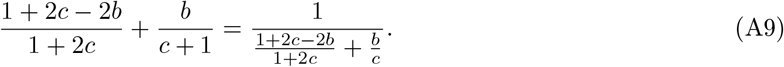

Solving for c and plugging back into (A8), we find

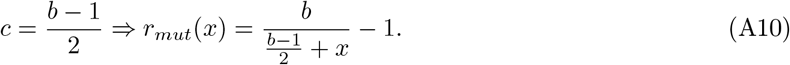

Finally, using the last condition in (A6), we can write

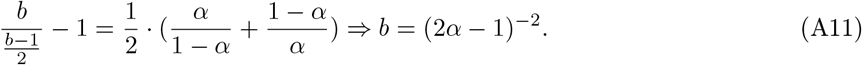

Therefore,

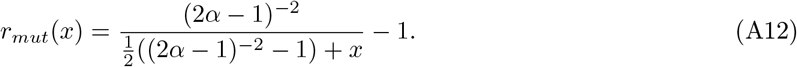

The mutant relative fitness can alternatively be written as

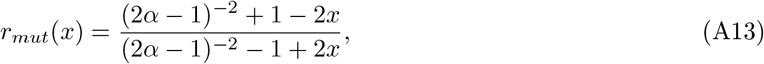

which more clearly establishes that this model is equivalent to an evolutionary game in which fitness is equal to payoff, with the payoff matrix

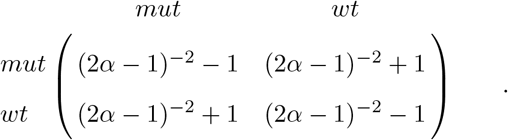

The overall mutant relative fitness is then

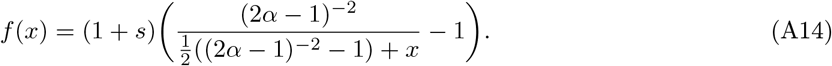

Note that at equilibrium, *f* (*x*) = 1. To find the equilibrium frequency *x*^*∗*^, we set *f*_*mut*_ = 1 and solve for x:

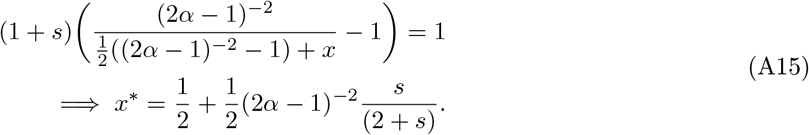

### A2: Deriving the fixation probability in the limit of the generalist population regime

In the limit of *α* ≈ 0.5, we can write

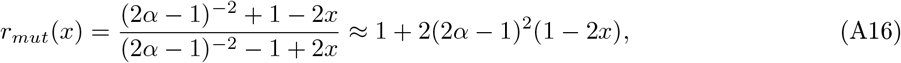

by Taylor expansion at *α* = 0.5. The total relative fitness can then be approximated as

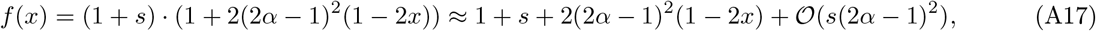

where we can ignore terms 𝒪 (*s*(2*α* − 1)^2^) since pure evolutionary selection is assumed to be weak throughout this paper. Similarly, the inverse eco-evolutionary fitness may be approximated as

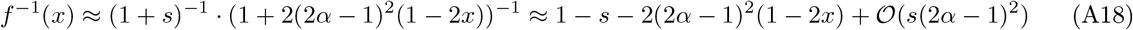

by Taylor expansion around *s* = 0 and *α* = 0.5.

The exact fixation probability in a well-mixed population is given by equation 1.16 in Traulsen and Hauert (2009) as

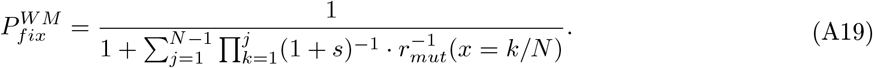

We can plug in *f* ^*−*1^ into (A31) to get

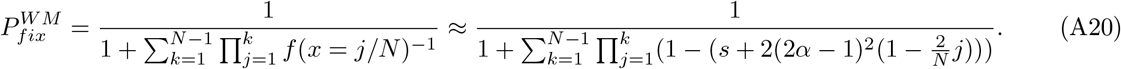

The product 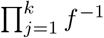 may be approximated as 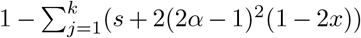, by retaining only the 𝒪 (*s*) and 𝒪 ((2*α* − 1)^2^) terms in the product. This simplifies to

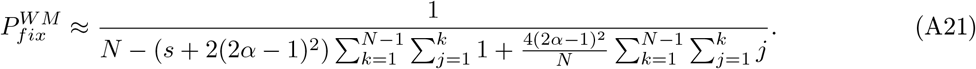

The two double sums can be approximated as

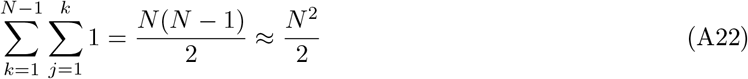

and

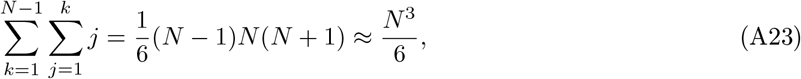

by retaining only leading order terms in N, assuming large N. Using a Taylor expansion around *s* = 0 and *α* = 0.5, we can write

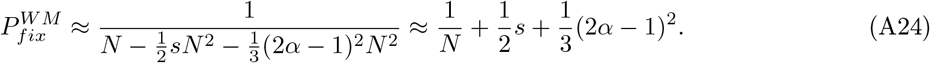

This approximation can be rewritten in terms of an effective selection coefficient as

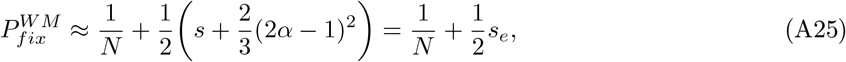

since the constant weak selection fixation probability has the form 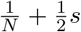 Allen, 2017)). It is helpful to write the fixation probability in this form because we can then capture the effect of spatial structure using *s*_*e*_ → *a*_*Bd*_ · *s*_*e*_.

### A3: Deriving approximations for *s*^*∗*^(*G*) and 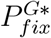 for large generalist populations

To find *s*^*∗*^(*G, α*), we first consider the difference in fixation probabilities and set it to 0

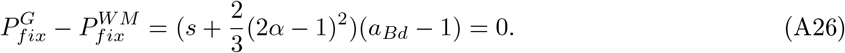

For a network G, the only solution to this equation is

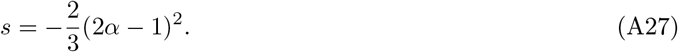

Therefore 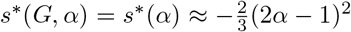 for generalist populations. To get 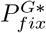, we plug *s*^*∗*^(*α*) into 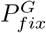 and can write

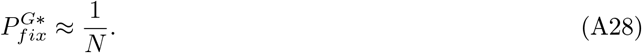

In the limit of large population size, 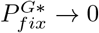.

### A4: Exact establishment and conditional fixation probabilities for specialist well-mixed and star-structured populations

The conditional fixation probability is the probability of mutant fixation starting from *Nx*^*∗*^ mutants. In specialist populations, *x*^*∗*^ = 0.5. The probability of mutant fixation given an initial mutant frequency of 0.5 can be computed using equation 1.20 in Traulsen and Hauert (2009) as

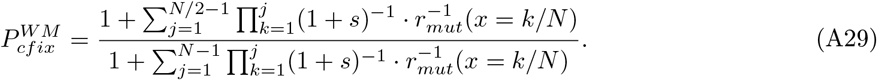

The establishment probability is then given by

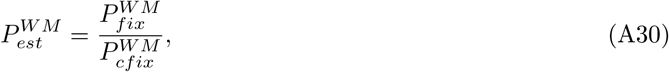

which follows from (9). The exact fixation probability on the star is given by equation 2.27 in Hadjichrysanthou et al. (2011) as

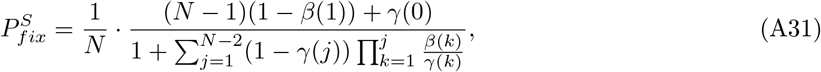

where

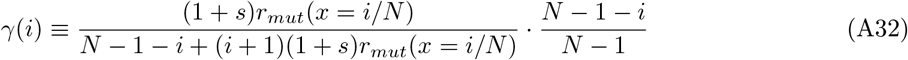

and

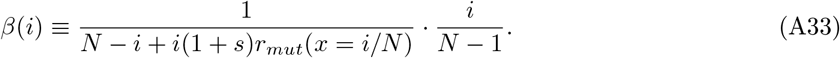

On the star graph, there are 2 possible configurations of *Nx*^*∗*^ mutants: 1) *Nx*^*∗*^ mutants on the leaf nodes and a wild-type on the center node or 2) *Nx*^*∗*^ − 1 mutants on the leaf nodes and a mutant on the center node. However, it is straightforward to see that configuration 1 can never be reached starting from a single mutant. Therefore the conditional fixation probability is equal to the probability of being absorbed in the state with N mutants starting from configuration 2. This is given by equation 2.23 in in Hadjichrysanthou et al. (2011):

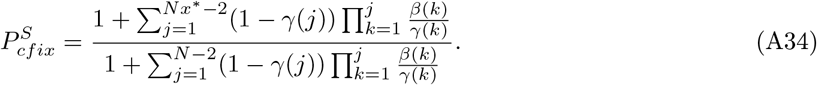

From (9), the exact establishment probability for a star can be computed by

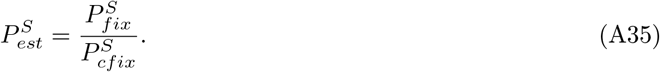

### A5: Deriving an approximation for the establishment probability for well-mixed specialist populations

The establishment probability in a well-mixed specialist population simplifies to

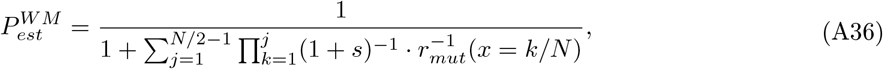

by evaluating (A30). Under strong ecological selection, the sum of products is dominated by the 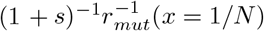 term, which may further be approximated as simply 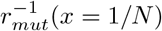, since pure selection *s* is weak. Therefore, by Taylor expansion around 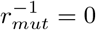 we write

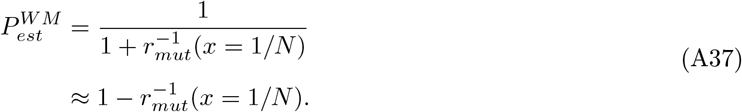

If we define the mutant invasion fitness as the fitness of the mutant when it has exactly one copy in the population, we can write

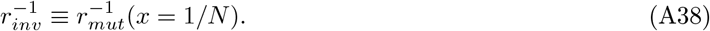

More explicitly, we can write

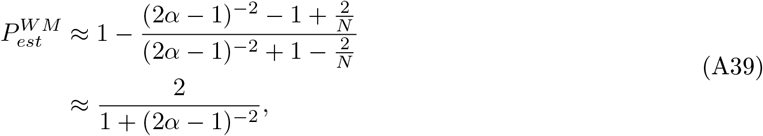

where in the last step we ignore 𝒪 (*N* ^*−*1^) terms.

### A6: Deriving an approximation for the conditional fixation probability for well-mixed specialist populations

Starting with the exact equation for the conditional fixation probability (A29), we can write

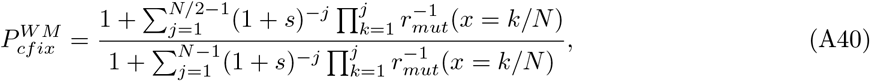

where we use 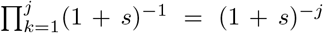. The key observation necessary to simplify 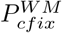 is that 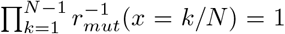, which follows from the symmetry property of *r*_*mut*_, i.e. 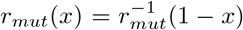.Therefore the last term in the sum in the denominator simplifies to 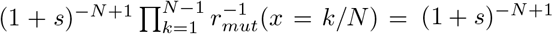. Pulling this term out of the sum, we can write

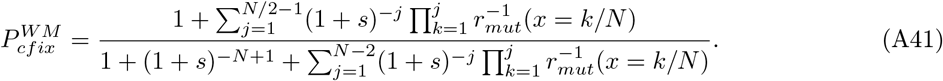

This term makes the largest contribution to the sum in the denominator. The next largest term in the denominator, and the largest in the numerator, is 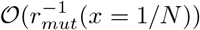. However 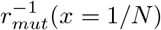 is very small under strong ecological selection, and therefore the contribution arising from 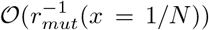 and all higher order terms may be ignored. This simplifies the problem to

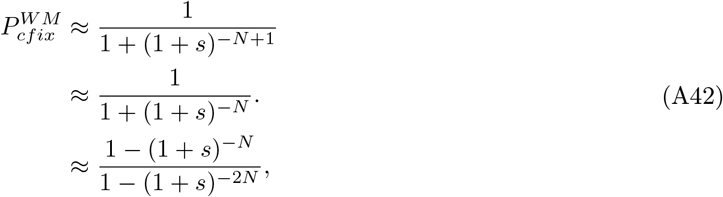

where, in the last step, we multiply both numerator and denominator by 1 − (1 + *s*)^*−N*^. We can alternatively write

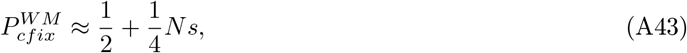

using Taylor expansion at *s* = 0.

### A7: Deriving approximations for *s*^*∗*^(*S, α*) and 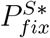, for large specialist populations

Using (A37) and (A43), we can approximate the total fixation probability in well-mixed specialist populations as

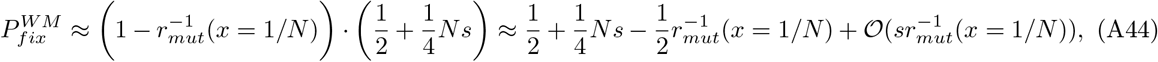

where we ignore 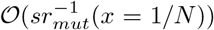 terms. On the star graph, s is amplified as *s* → 2*s* (Kuo et al., 2021) and *r*_*mut*_(*x* = 1*/N*) as 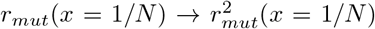 (Lieberman et al., 2005), because s is weak and *r*_*mut*_(*x* = 1*/N*) is strong. Therefore, the fixation probability on the star can be approximated as

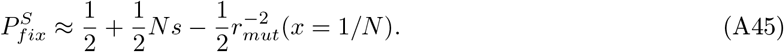

To understand the point at which the fixation probabilities in the well-mixed and star populations intersect, we take the difference set to 0,

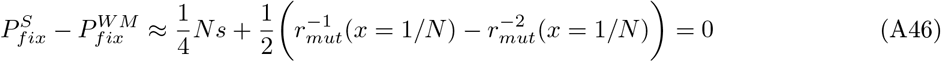

and solve for *Ns*^*∗*^, which allows us to write

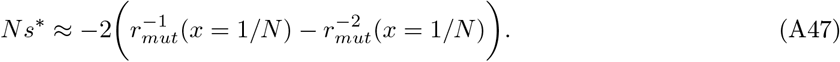

This equation showcases the value of *Ns* at which the fixation probabilities intersect for any given (strong) *α*. Since *r*_*mut*_(*x* = 1*/N*) is large in the specialist regime, we can drop the 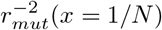 term and write

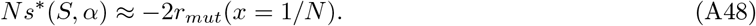

Plugging 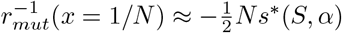 into 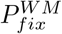 gives us 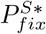

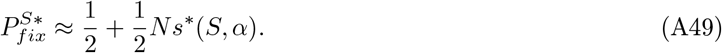

## Supplementary Figures

**Figure S1:**
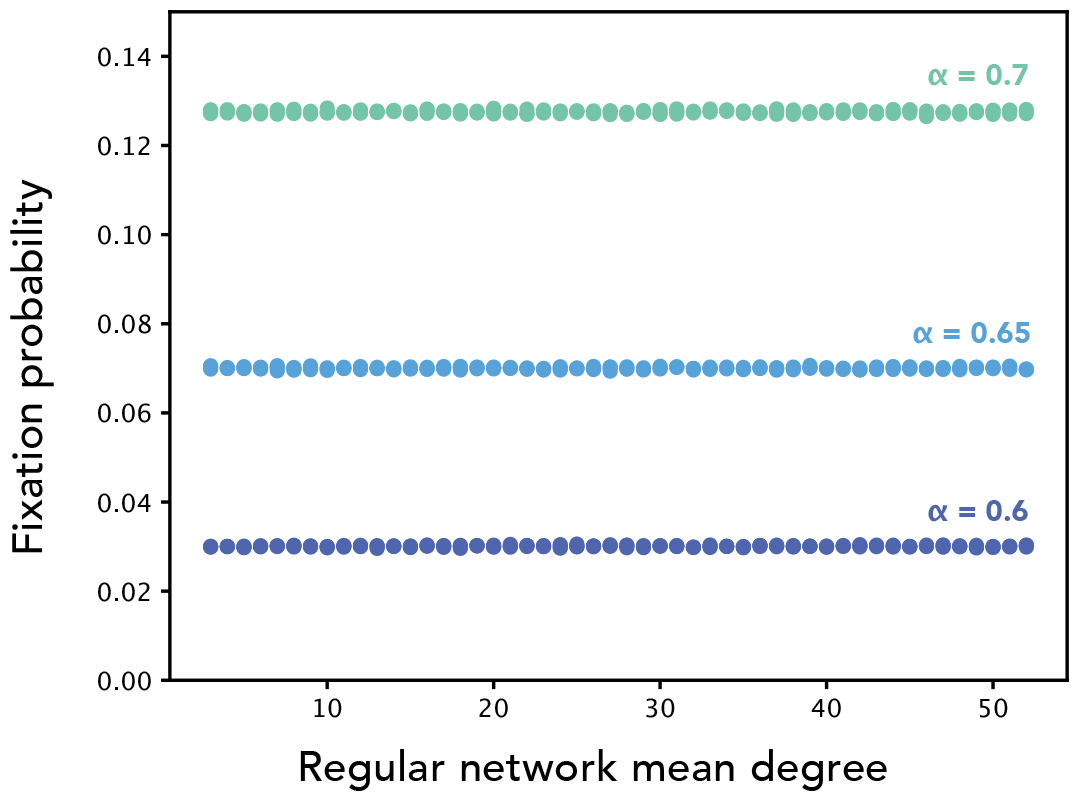
The isothermal theorem holds under global frequency-dependent dynamics. Each point shows the fixation probability of the mutant ecotype on a k-regular graph with mean degree as on the x-axis and N=100 for various values of *α*. Here *s* = 0. Fixation probabilities are computed using simulations with at least 2e6 replicates.

**Figure S2:**
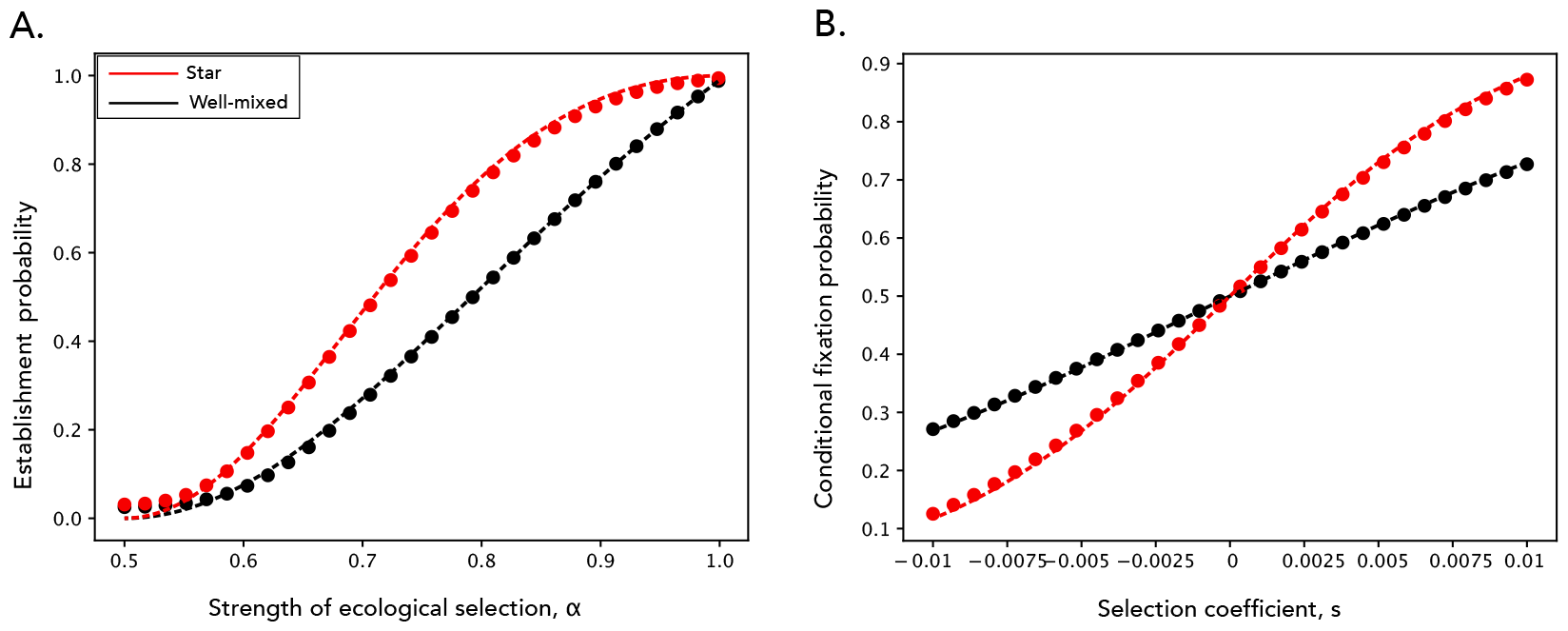
Approximations for establishment and conditional fixation probabilities in well-mixed and star-structured populations match exact calculations. **Panel A** Dots show exact establishment probabilities on the complete and star graphs for varying ecological selection strength *α* and fixed pure selection strength *s* = 0.01. Solid lines show the analytic approximations. Here *N* = 100. **Panel B** Dots show exact conditional fixation probabilities on the complete and star graphs for varying pure selection strength *s* and fixed ecological selection strength *α* = 0.9. Solid lines show the analytic approximations. Here *N* = 100.

**Figure S3:**
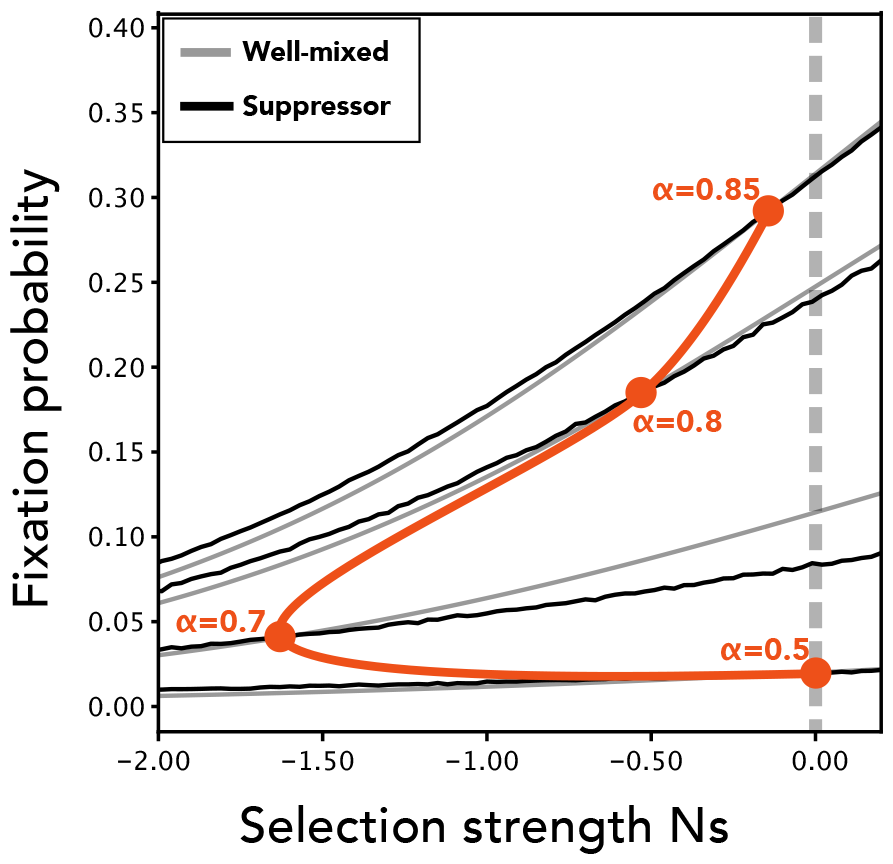
Ecological interactions can reverse the role of amplifiers and suppressors for weakly deleterious mutants. The case of a suppressor topology. The black lines are fixation probabilities estimated through 10^6^ simulation runs for a suppressor network (amplification factor 0.47), for four different ecological strengths (values as depicted next to the orange dots). The grey lines shows the corresponding, exact fixation probabilities for a well-mixed population. The orange line follows the point of intersection between the well-mixed and suppressor populations, as ecological strength *α* is continuously varied. Population size *N* = 50.

**Figure S4:**
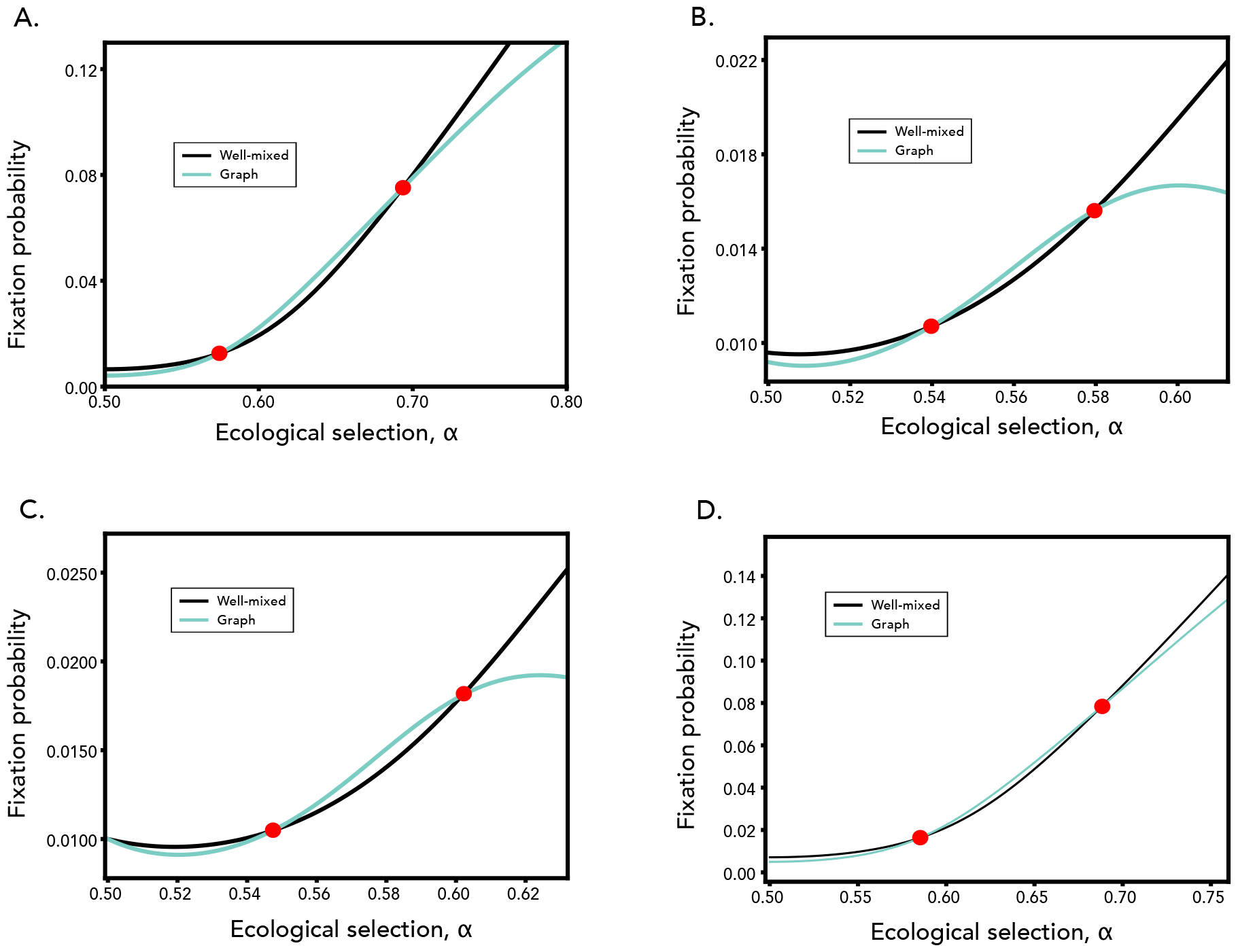
Robustness to asymmetric metabolic profiles and supply rates. **Panel A**. Fixation probabilities of a well-mixed (black) and star (green) population. Here, population size *N* = 100 and *s* = − 0.008. **Panel B. Fixation probabilities under different metabolic profiles for each variant**. We assume the wild-type has metabolic profile [*α*, 1 − *α*] and the mutant [1 − (*α* − *δ*), *α* − *δ*]. We set *δ* = − 0.025. The mutant has a pure deleterious selection coefficient because it’s metabolic profile is not as well matched to the supply rate as the wild-type. **Panel C. Fixation probabilities under different supply rates for each resource**. We assume that the mutant’s preferred resource is supplied at rate 0.5 + *δ* and the other resource at 0.5 − *δ*. We set *δ* = − 0.025. **Panel D. Fixation probabilities under deleterious pure selection coefficient and structured resources**. Here, *N* = 100 and *s* = − 0.012. All resources are externally supplied at an equal rate and depleted at rates that vary spatially, determined by the identity of individual occupying that space. Resources are allowed to diffuse along the edges of the network with diffusion coefficient *D* = 2.

**Figure S5:**
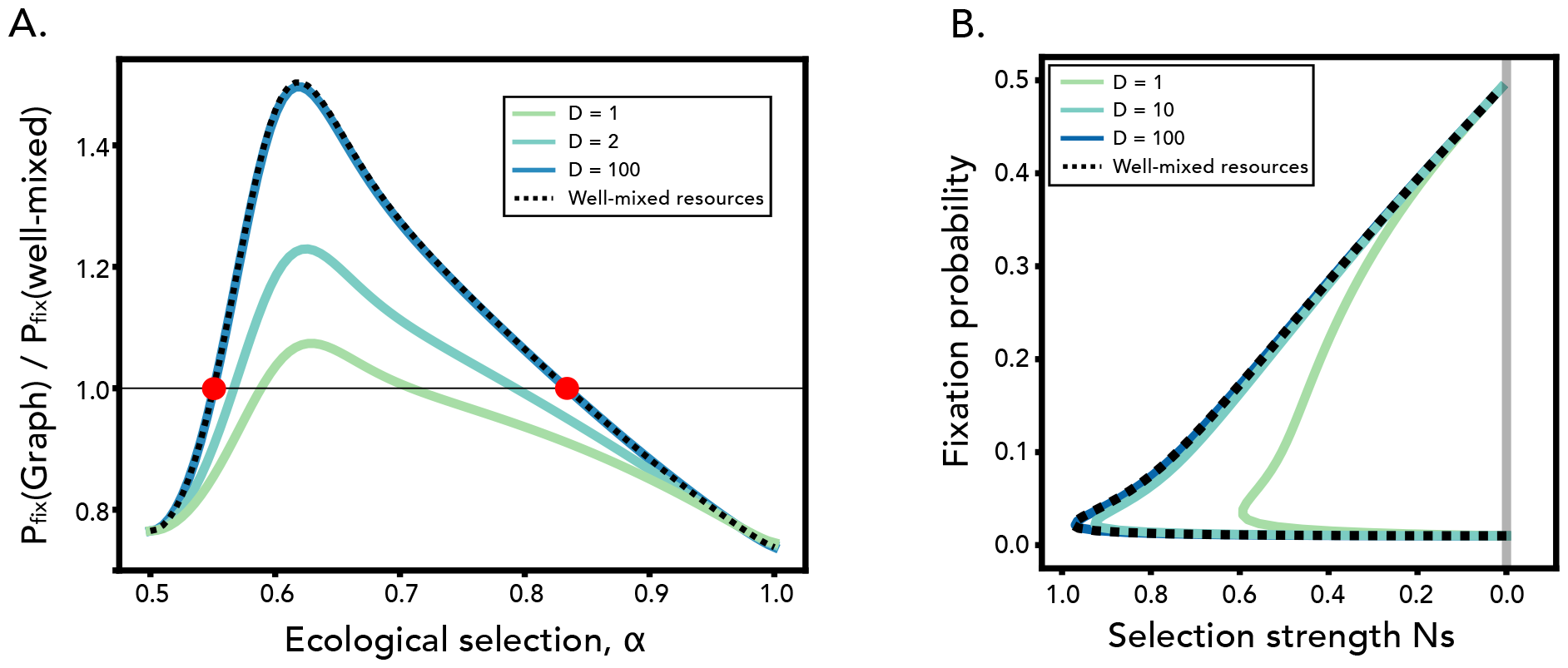
Robustness of our results to diffusible resources. In this model, all resources are externally supplied at an equal rate and depleted at rates that vary spatially, determined by the identity of the individual occupying that space. Resources are allowed to diffuse along the edges of the network with diffusion coefficient D. This coefficient is varied as in the legend. **Panel A**. Fixation probability ratios for a population with *N* = 100. Pure selection coefficient *s* = − 0.005. **Panel B**. Points of intersection between fixation probabilities between the network and well-mixed model for various diffusion coefficients, same parameters as in Panel A.

## Notes

### Competing Interest Statement

The authors have declared no competing interest.

